# A 39.8kb flavi-like virus uses a novel strategy for overcoming the RNA virus error threshold

**DOI:** 10.1101/2024.01.08.574764

**Authors:** Mary E. Petrone, Joe Grove, Jonathon C. O. Mifsud, Rhys H. Parry, Ezequiel M. Marzinelli, Edward C. Holmes

## Abstract

It is commonly held that there is a fundamental relationship between genome size and error rate, manifest as a notional “error threshold” that sets an upper limit on genome sizes. The genome sizes of RNA viruses, which have intrinsically high mutation rates due to a lack of mechanisms for error correction, must therefore be small to avoid accumulating an excessive number of deleterious mutations that will ultimately lead to population extinction. The proposed exceptions to this evolutionary rule are RNA viruses from the order *Nidovirales* (such as coronaviruses) that encode an error correcting exonuclease, enabling them to reach genome lengths greater than 40kb. The recent discovery of large genome flavi-like viruses (*Flaviviridae*), which comprise genomes up to 27kb in length yet seemingly do not encode exonuclease domains, has led to the proposal that a proofreading mechanism is required to facilitate the expansion of RNA virus genomes above 30kb. Herein, we describe a 39.8kb flavi-like virus identified in a *Haliclona* sponge metatranscriptome that does not encode an exonuclease. Structural analysis revealed that this virus may have instead captured bacterial domains associated with nucleic acid metabolism that have not been previously found in RNA viruses. Phylogenetic analysis placed this virus as a divergent pesti-like lineage, such that we have provisionally termed it Maximus pesti-like virus. This virus represents the first instance of a flavi-like virus achieving a genome size comparable to that of the *Nidovirales* and demonstrates that RNA viruses have evolved multiple solutions to overcome the error threshold.

## INTRODUCTION

RNA viruses have traditionally been characterised by small, compact genomes, often close to 10 kb in length^1^. The longest RNA virus genome documented to date is ~47kb^2^, considerably shorter than those seen in DNA viruses whose genomes can exceed 2.5Mb^3^. Despite the explosion in RNA virus discovery following the advent of metagenomic sequencing^4-7^, the maximum genome sizes of most families of RNA viruses have remained relatively stable, with the longest genomes (i.e., >30 kb) consistently falling within families from the order *Nidovirales*.

The most popular theory for the restricted genome sizes of RNA viruses is that this stems from the mutational burden associated with replication via an error-prone RNA polymerase (i.e., RNA dependent RNA polymerase (RdRp) or reverse transcriptase)^8^. Because these enzymes usually lack any form of error correction such as proofreading, longer genomes would be expected to accumulate more mutations per genome replication. As most mutations are deleterious (or even lethal)^9^, a high error rate will greatly reduce virus fitness, eventually leading to population extinction. Consequently, an upper cap on genome sizes reflects an “error threshold”, with high mutation rates necessarily resulting in smaller genomes.

The existence of a mutation-driven cap on virus genome size is supported by two lines of evidence. First, the treatment of experimental populations of RNA viruses with mutagenic agents, such as 5-azacitydine and 5-fluorouracy, results in major fitness losses as expected if these viruses reside close to the error threshold^10^. Indeed, fitness losses resulting from the use of antiviral mutagenic agents such as ribavirin and molnupiravir^11^ underpins viral therapies based on the induction of so-called “lethal mutagenesis”^12,13^. Second, members of the *Nidovirales*, which include those RNA viruses with the largest known genomes^2^, have acquired a unique proofreading mechanism. The *Coronaviridae* encode endo- and exoribonucleases (such as nsp14-ExoN) that are essential for replication^14^ and that correct replication errors^15,16^. Similarly, members of the *Roniviridae* utilise NAD and ADP-ribose (NADAR) domains which are associated with RNA repair^17^. Reducing the error rate per nucleotide should, in theory, have enabled nidovirus genomes to expand in size. Indeed, the discovery of Nam Dinh virus, a ~20kb nidovirus that is distantly related to the *Coronaviridae* and the *Roniviridae* and encodes an exoribonuclease further supports this hypothesis^18^.

Although attractive, the error threshold theory for virus genome size was challenged by the discovery of the so-called “large genome flavi-like” viruses (LGFs) that possess genomes up to 27kb in length^19,20^. These place the LGFs as the second longest RNA virus lineage documented to date; importantly, however, they do not encode an exoribonuclease or similar domain. As a result, it was proposed that a threshold of 30kb in fact represented the upper limit for genome size in RNA viruses in the absence of proofreading, provided that other factors such as a stabilising methyltransferase domain were present^21^.

Herein, we describe a novel, flavi-like virus with a genome of ~40 kb that does not encode a recognisable proofreading domain. This discovery provides new insights into the evolution of the *Flaviviridae* and demonstrates that exonuclease domains are not required for RNA virus genome expansion above 30kb, challenging assumptions about the factors that determine RNA virus genome size.

## RESULTS

### Discovery of a large and highly divergent marine flavi-like virus

We identified a 39.8kb flavi-like virus in a sea sponge (order Haplosclerida) metatranscriptome. This sponge, along with 72 invertebrate specimens, was collected from Chowder Bay, Sydney Harbour, Sydney, Australia on October 20, 2022. Assessment of the true host association of this virus was challenging due to the complexity of the sequencing library (**Supp. Fig. 1a**), which in turn reflected the heterogeneous composition of the ecosystem from which it was collected (**Supp. Fig 1b**). Despite this, over 80% of rRNA reads in this library shared high sequence similarity with rRNA of sponge, particularly those belonging to the genus *Haliclona*, order Haplosclerida (**Supp. Data**). Given that the morphology of the specimen was consistent with *Haliclona* (**Supp. Fig. 1b, *inset***) and this genus has been previously documented in Sydney Harbour (ozcam.ala.org.au), we concluded that a *Haliclona* sponge was the most probable host organism of this novel virus.

This virus exhibited clear, yet minimal, sequence similarity with members of the *Flaviviridae*. It was most closely related to Fushun flavivirus 1 (length = 15kb), with which it shared 25.7% sequence similarity (e-value = 6.36e-25). Like other *Flaviviridae*, it comprised a single polyprotein (38kb) and long 5’ and 3’ untranslated regions (UTRs) (462 nt and 1282 nt, respectively) (**Fig. 1a**). The polyprotein sequence was complete, containing no premature stop codons and exhibited a strong AT bias (64%). Its abundance in the sequencing library was relatively low, comprising only 0.11% of non-rRNA reads (58,520 reads mapped to the flavi-like virus contig), but its sequencing coverage was robust, with a mean depth of 210.7 (st. dev. = 86.3) (**Fig. 1a**). Fragments of this virus were detected in two of the three other libraries of samples collected from the same piling (Bryozoa, 0.0001% non-rRNA reads; sponge, 0.007% non-rRNA reads) (**Supp. Fig. 2**^22^). To our knowledge, this is the longest known flavi-like virus, nearly 1.5 times the length of previously documented LGFs. It is also the longest known RNA virus outside of the order *Nidovirales* and the only known RNA virus over 30kb to encode a single polyprotein.

**Figure 1:**
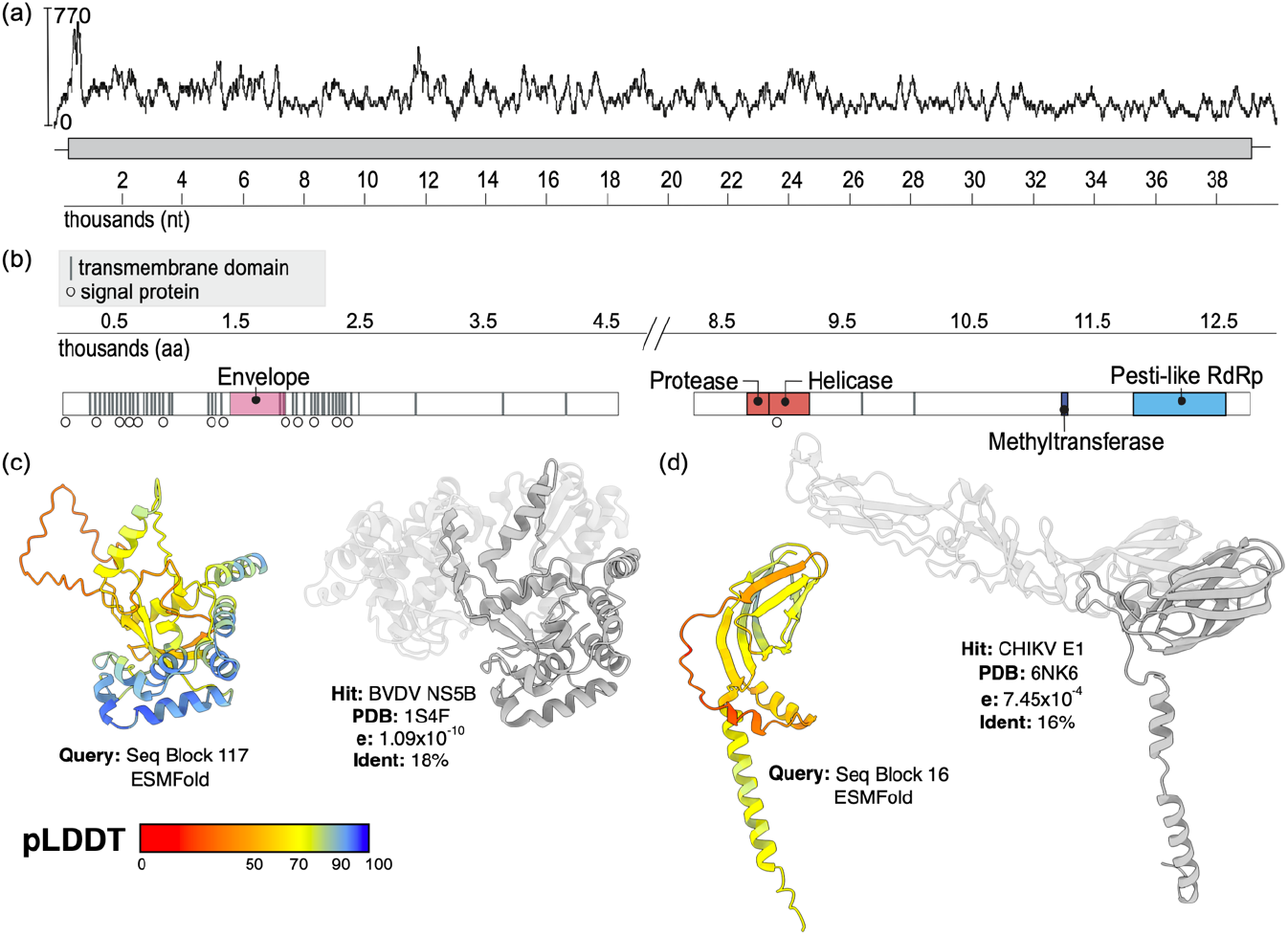
Characterisation of canonical *Flaviviridae* features in a highly divergent, large genome flavi-like virus. (a) Sequencing coverage of nucleotide sequence. The y-axis measures sequencing depth. Reads were mapped using BBMap^26^ and the coverage plot was generated in Geneious Prime. (b) Functional domains identified using sequence-based and protein fold-based homology. (c) Structural prediction of the putative NS5 gene. Left: ESMFold structure prediction for polyprotein sequence block 117 (amino acid residues 11700-12000), color-coded by prediction confidence (predicted Local Distance Difference Test, pLDDT) as shown in the key. Right: homologous BVDV NS5B structure (PDB:1S4F) identified by FoldSeek. Expected value and percentage sequence identity are shown. Regions of structure not aligned by FoldSeek are transparent. (d) Structural prediction of putative glycoprotein. Homology between sequence block 16 and Chikungunya virus (CHIKV) E1 structure (PDB: 6NK6), displayed as in (c).

Despite its length and low sequence similarity to the *Flaviviridae*, this virus possessed characteristics of canonical flavivirids. To explore genome organisation and annotate protein function we used both sequence-based approaches (BLASTp, InterProScan, SuperFamily, NCBI CD-Search, and NCBIfam) and protein structure prediction followed by fold-based homology searches. For the latter the polyprotein sequence was broken into overlapping 300 residue blocks for structure prediction, using AlphaFold^23^ and ESMFold^24^, and queried using the FoldSeek^25^ server. The putative NS2/3 gene featured a consecutive serine protease and helicase pair including a helicase domain (**Fig. 1b**). Manual annotation showed that the serine protease included the typical Ser (nucleophile), His (base), Asp (acid residue) catalytic triad. Similarly, the putative NS5 gene was pesti-like in both sequence and structure (**Fig. 1c**) and retained the highly conserved GDD motif. An envelope protein surrounded by signal peptides and probable N-linked glycosylation sites was detected near the 5’ terminal (**Fig. 1b,d, Supp. Fig. 3**). ESMFold protein structure prediction yielded a domain of intermediate prediction confidence towards the C-terminus of this region. This domain exhibited similarity to domain III of a class II fusion protein, with strongest homology to glycoproteins from bunya- (Gc) and alphaviruses (E1) rather than the E protein found in orthoflaviviruses (Phyre2: Rift Valley Fever virus, HHPred: Thrombocytopenia syndrome virus, FoldSeek: Chikungunya virus). This is consistent with the highly divergent nature of this novel protein, as this region exhibited only 16% identity with its closest homolog (**Fig. 1d**). Like other LGFs, a methyltransferase domain was present within the putative NS5. However, we found no regions that were similar to nido-like endo/exonucleases or NADAR domains (**Fig. 1b**). The quality of the sequence and the organisation of the putative genome suggested that this unusually large virus was exogenous and related, although distantly, to the *Flaviviridae*.

### Divergent, bacteria-associated domains in an RNA virus genome

Given its size, we hypothesised that this virus would encode non-canonical flavivirus domains, particularly those that would help support the replication of such a large RNA genome. Accordingly, using both sequence- and structure-based approaches, we found three notable features: a novel extended helical domain (pos. 4,346-4,475), a Tu elongation factor (pos. 6,419-6,586), and a nucleic acid metabolism cassette (pos. 7,286-8,206) (**Fig. 2a**).

**Figure 2:**
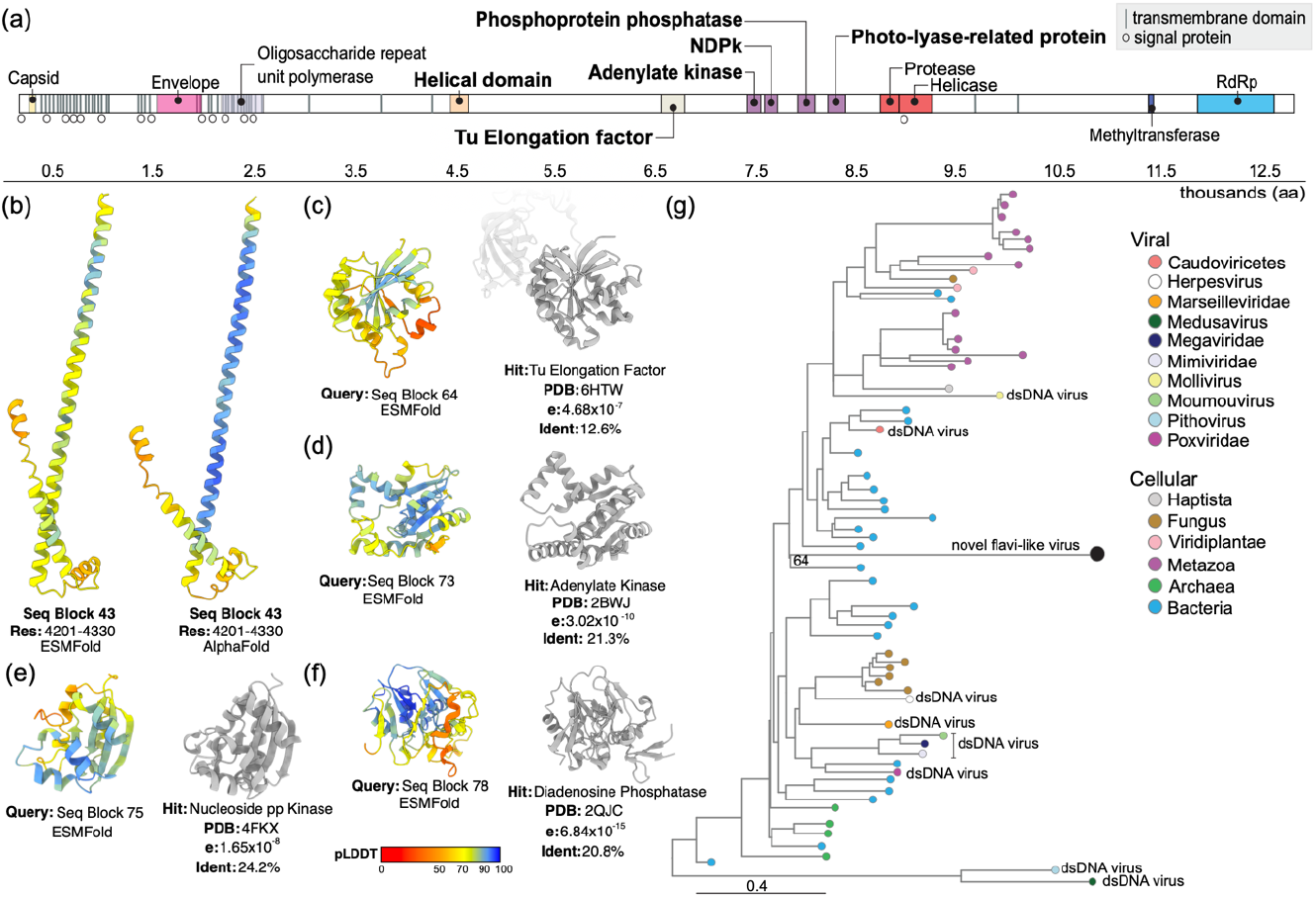
Large genome flavi-like virus polyprotein encodes a complement of non-canonical functional domains. (a) Annotation of domains detected using a combination of sequence and structure-based analyses: InterProScan, DeepTMHMM (transmembrane domains), SignalP (signal peptidases), Pfam, CDD, and FoldSeek. Domain annotations reflect domains sharing the closest structural homology to the putative viral domains. The polyprotein is scaled to thousand amino acids. NDPk: nucleoside diphosphate kinase. Helical domain: helical domain of unknown function. (b) Inferred ESMFold and AlphaFold structures of a helical domain of unknown function (found in sequence block 43), color-coded by prediction confidence (pLDDT) as shown in the key (c-f) ESMFold structures of the stated sequence blocks alongside their cognate FoldSeek hits, expected value and percentage sequence identity are provided. Regions of structure not aligned by FoldSeek are transparent. (g) Midpoint rooted maximum likelihood phylogenetic tree of NDPk domains. Tip colours indicate the organism associated with each sequence. Ultrafast bootstrap values are shown as numbers at select nodes. The tree branches are scaled according to the number of amino acid substitutions per site.

The potential functions of these domains were uncertain. The role of the extended helical domain could not be determined. Both ESMFold and AlphaFold structures were of intermediate to high confidence and were in excellent agreement, supporting their veracity. However, this feature did not share detectable homology with known proteins in both ESMFold and AlphaFold predictions (**Fig. 2b**). Its position in the genome proximally downstream of the envelope protein could be consistent with host response antagonism like the canonical NS1^27^, although experimental analysis is required to test this. In cellular organisms, Tu elongation factors (**Fig. 2c**) principally facilitate translation^28^. This may be particularly beneficial to a virus with a very long polyprotein, but Tu elongation factors are also thought to exhibit ‘moonlighting’ (i.e., non-canonical) functions^28^. Moreover, in this instance the protein was highly divergent, exhibiting no detectable sequence similarity to any other protein in the NCBI nr database, including from cellular organisms. This may reflect functional divergence. We identified related domains in the genomes of some *Megaviricetes* and *Caudoviricetes* (bacteriophage) when we screened the Reference Proteomes database with HMMER^29^, although we found no homologues in RNA viruses. The predicted structure shared homology with eukaryotic and bacterial elongation factors, but it was most closely related (12.6% identity) to a Tu Elongation Factor characterised in *Campylobacter jejuni* (Gram-negative bacteria) (**Fig. 2c**).

Elements of the nucleic acid metabolism cassette were more conserved. This region comprised an adenylate kinase, a nucleoside diphosphate kinase (NDPk), a phosphoprotein phosphatase, and a photo-lyase-related protein upstream of the NS2/3 protease-helicase pair (**Fig. 2a,d-f**). Among these domains, NDPks are the most highly conserved across all kingdoms of life^30^ and have been found in some giant DNA viruses and bacteriophage^31^. They are involved in RNA synthesis but have not previously been documented in RNA viruses. Interestingly, the flavi-associated NDPk domain was more closely related to those in cellular organisms than to those in other viruses, sharing up to 30% sequence similarity to homologous domains in bacteria and eukaryotes. To search for related domains in viruses, we screened the NCBI non-redundant (nr) protein database restricted to viruses (taxid: 10239) using the flavi-associated NDPk as input. This returned no results, suggesting that the flavi-associated NDPk domain was not related to its counterparts in DNA viruses. It also suggested that NDPk acquisition is a rare event in RNA virus evolution. Querying Reference Proteomes, UniProtKB, Swiss-Prot, PDB, and AlphaFold databases using hmmsearch with the NDPk alignment as input also failed to detect this domain in any publicly available RNA virus genomes.

Phylogenetic analysis of the NDPk domain further supported the hypothesis that the flavi-associated domain was derived from bacteria. Specifically, this sequence fell in a clade with the NDPk of *Candidatus Marinimicrobia* bacterium, albeit with moderate support (ufboot = 64) and a long branch (**Fig. 2g**). These features may be indicative of the differing evolutionary rates in cellular and viral organisms. Interestingly, the NDPk domains from DNA viruses did not form a monophyletic group suggesting there have been multiple acquisition events from either fungi or bacteria. The inferred structure of the flavi-associated domain also suggested a closer relationship with bacterial NDPk (**Fig. 2e**). These observations support the conclusion that this cassette was likely acquired from bacteria, constituting the only known example of such a feature in the characterised RNA virosphere.

### Phylogenetic support for classification as a large genome flavi-like virus

The unique and divergent protein complement encoded by this virus suggested that it could provide new insights into the evolutionary history of the *Flaviviridae*. A family-level phylogenetic analysis placed both the putative NS2/3 and NS5 genes with pesti-like viruses, albeit in a deep phylogenetic position and without robust support (**Fig. 3a,b**). We herein refer to this section of the tree as the *Pestivirus*-LGF clade, which comprises the genus *Pestivirus*, pesti-like viruses, and LGFs. The placement of the novel NS5 among LGFs was moderately stable (ufboot = 72), and the size of the viral genome was consistent with other members of this group (**Fig. 3a, *inset***). In contrast, the putative NS2/3 fell towards the base of the clade containing canonical pestiviruses with moderate support (ufboot = 79) (**Fig. 3b, *inset***). Previously, two spider-associated pest-like viruses (Xinzhou spider virus 3 and Shayang spider virus 4) were the most divergent members of this clade^32^.

**Figure 3:**
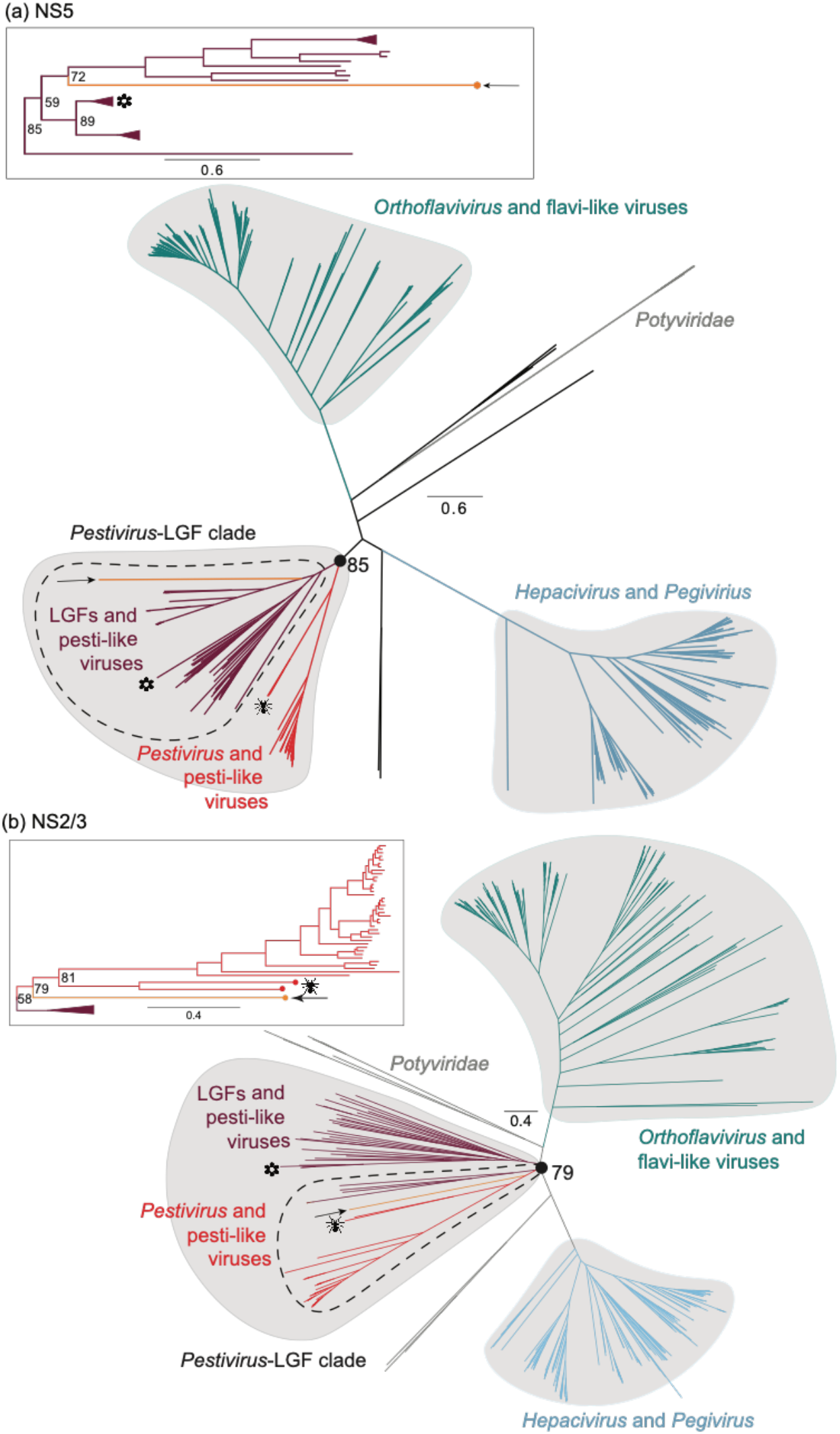
Phylogenetic analysis of the novel flavi-like virus places it among the pesti-like viruses. Unrooted phylogenetic trees of *Flaviviridae* (a) NS5 and (b) NS2/3. Arrows indicate the novel flavi-like virus identified here. Flower icon indicates plant-associated flavi-like viruses. Spider icon indicates spider-associated pesti-like viruses. Ultrafast bootstrap values are shown as numbers at select nodes. Insets show the groups containing the novel flavi-like virus and correspond to the regions encircled by dotted lines in the main figure. The topology of the insets reflects midpoint rooting of each tree in the main figures. Tree branches are scaled according to amino acid substitutions.

In an attempt to improve the accuracy of the phylogenetic position in both genes we realigned all NS2/3 and NS5 sequences belonging to the orthoflavivirus, pestivirus, pesti-like viruses, and LGFs using six combinations of sequence alignment and alignment trimming methods (**Table 1**). This did not yield more stable results for the placement of either region (**Supp. Fig. 4,5**). However, both gene sequences from the virus newly identified here consistently fell at the base the *Pestivirus*-LGF clade, suggesting that our novel virus represents part of the deep evolutionary history of this group. Both midpoint rooting and rooting on the *Orthoflavivirus* clade consistently placed our novel virus within the sister clade to that containing the genus *Pestivirus* and without a close phylogenetic relationship to known plant- or protist-associated pesti-like viruses. If virus-host codivergence is assumed, the placement of this virus deep in the phylogeny is consistent with an ancient invertebrate host. This was particularly true for the novel NS2/3 sequence, which formed a sister group to the spider-associated pesti-like viruses. Overall, these analyses suggest that this virus is most likely a large genome sponge-infecting pesti-like virus, and as such we have provisionally named it Maximus pesti-like virus.

**Table 1.**
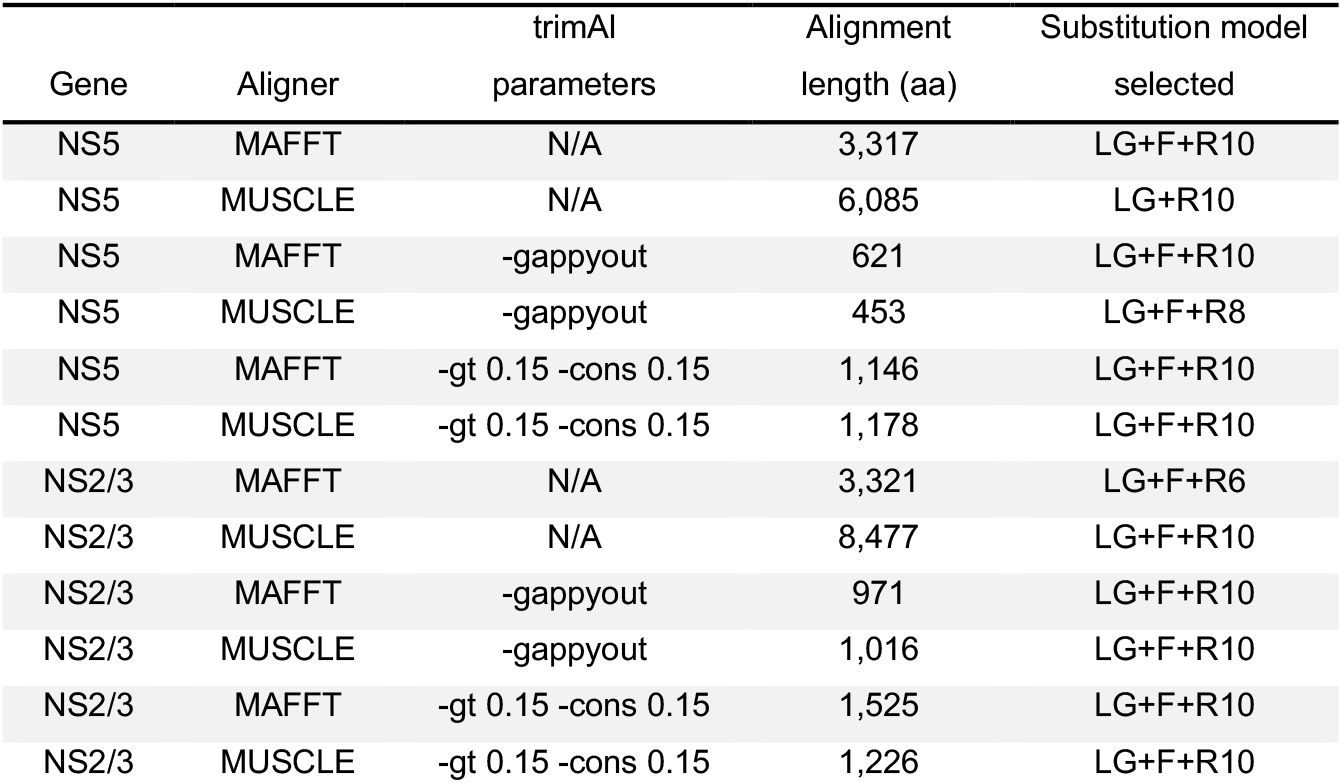
Combinations of sequence alignment and trimming methods used in phylogenetic analysis of the *Pestivirus*-LGF clade.

## DISCUSSION

We describe the largest known flavi-like virus and largest known RNA virus outside of the order *Nidovirales*. With a genome size of 39.8kb, this virus surpassed the longest documented flavi-like virus to date by 12kb^19^. Despite its large size, this virus possessed features that are characteristic of members of the *Flaviviridae*: an E-like glycoprotein, a protease/helicase pair, and a pesti-like RdRp. Although the E-like glycoprotein shared highest similarity to alpha- and bunyavirus glycoproteins, this is consistent with the notion that the class II membrane fusion machinery of the *Amarillovirales* (that contain the *Flaviviridae*), *Bunyavirales*, and *Martellivirales* evolved from a common ancestor^33^. The polyprotein encoded a methyltransferase upstream of the RdRp, which is common throughout the *Flaviviridae*^21^. The organisation of the genetic components was also congruent with that of canonical *Flaviviridae*, a characteristic that differs from large nidoviruses, as these exhibit a flexible modularity in their genetic organisation^2^. Phylogenetic analysis placed the two most conserved genes (NS2/3 and NS5) of this novel virus deep within the *Pestivirus*-LGF clade, with a relatively close relationship to previously characterised LGFs. Although small portions of the genome shared sequence and structural homology to cellular genes, we concluded that the virus was unlikely to be an endogenous viral element because the virus-associated sequences were highly divergent from their cellular homologues.

The discovery of such a large flavi-like virus raises important questions. Does it possess a mechanism for correcting errors during replication? Or does it sustain a mutation rate similar to those seen in other flavivirids, and if so, how? Importantly, we did not find evidence for nido-like endo- or exonucleases, the absence of which is consistent with other LGFs. Instead, the nucleic acid metabolism cassette encoded upstream of the putative NS2/3 could be a plausible proofreading candidate. One component of this cassette, deoxyribodipyrimidine photo-lyase is involved with nucleotide repair caused by UV damage^34^. However, other components may serve different purposes. Adenylate kinases regulate adenosine phosphate concentrations^35^, and, when present in dsDNA viruses, are thought to counterbalance the differential AT concentrations between the virus and its host along with NDPk^31^. The genome of our novel flavi-like virus exhibited a substantial AT bias (64%), and the nucleic acid metabolism cassette may in part function to sustain this. The lack of additional proofreading candidates reflected our inability to annotate or assess the role of large portions of the genome *in silico* because they did not share sequence or structural homology with known proteins. Isolation and culturing of this virus is needed to ascertain the function of these proteins. *In vitro* studies are also needed to estimate the intrinsic mutation rate of this virus so that it may be compared to those of canonical, 10kb vertebrate flaviviruses (e.g., hepatitis C virus, dengue virus).

The genomic architecture of this virus may also be consequential in understanding the long-term evolutionary history of the *Flaviviridae*. In contrast to nidoviruses, which are thought to have undergone genomic expansion after the acquisition of a proofreading mechanism, members of the *Pestivirus*-LGF clade may have undergone genome reduction. The phylogenetic placement of NS5 sequences within the *Pestivirus*-LGF clade was uncertain, but this uncertainty was mitigated by the consistent placement of the novel NS5 across multiple combinations of sequence alignment and alignment trimming methods, and the relatively strong support for this clade as a whole (ufboot = 85) in the family-level phylogeny. The position of the NS2/3 gene was better supported (ufboot = 79) near the base of the clade containing canonical pestiviruses. This tree topology would be consistent with the phylogenetic position of its putative host, a sponge, which was among the first animals to evolve^36^. Thus, one interpretation of the size of the Maximus pesti-like virus genome is that it is a relic of an ancestral state of the pestiviruses, and the canonical (i.e., 10kb) *Pestivirus* genome resulted from a series of genome reduction events, perhaps concordant with their emergence in vertebrates where longer genomes would mean more potential immune targets for organisms that had evolved adaptive immunity^37^. If large sections of the *Pestivirus* genome were lost over time, it follows that the proteins encoded in these sections were selected against. This could explain why we were unable to find similar protein domains in other RNA viruses. It also raises the interesting possibility that some RNA viruses previously encoded a proofreading mechanism and then lost it, if one is found in the novel flavi-like virus. Such a pattern would again be opposite to what has been inferred from the evolutionary history of the *Nidovirales*.

Another explanation for the aberrant size of this novel virus is that it belongs to a select group of specialised viruses that are successful “genetic pirates”. Although its conserved regions fall within the *Pestivirus*-LGF clade, large sections of its genome could not be aligned to any known member of the *Flaviviridae* and were not present in any RNA virus searchable in publicly available databases. Thus, the acquisition of the nucleic acid metabolism cassette and the other functional domains could have been the result of relatively recent genetic piracy wherein an ancestral virus captured genetic material from a cellular organism. The novel flavi-like virus appears to have acquired these genes in at least two separate events because its putative elongation factor does not share detectable sequence similarity with known cellular homologues, whereas its NDPk domain does. This suggests that the ancestors of this virus may have been adept at stealing genes from cellular organisms, a characteristic that may not be representative of most ancestral *Flaviviridae*. A more confident assessment of these scenarios will require the discovery of additional LGFs of comparable size.

Regardless of when it occurred, the apparent acquisition of bacterial genes is puzzling. The phylogenetic placement of the flavi-associated NDPk domain suggests that it was acquired directly from bacteria (**Fig. 2c**). Bacterial NADAR and Nudix domains have been found in RNA viruses, including among the *Nidovirales*, but these are thought to have been acquired through horizontal gene transfer from a eukaryote^17,38^. A parsimonious interpretation of our NDPk phylogenetic tree is inconsistent with this sequence of events because the flavi-associated NDPk sits with bacteria-associated NDPk domains. One explanation is that ancestral viruses infected bacteria-eating protists. Because bacteria have evolved defences that allow them to use their would-be predators as safe ecological niches^39^, co-infection with an ancestral flavi-like virus could have afforded the virus an opportunity to steal bacterial genes. Alternatively, sponges harbour many endosymbionts, including bacteria^40^, which may have facilitated bacteria-virus gene transfer. It may also be that the flavi-associated NDPk shares closer sequence similarity to an unsampled eukaryote, and its position in the phylogenetic tree will change as new genetic data become available.

Filling sampling gaps in the RNA virosphere may shift paradigms in evolutionary virology if more exceptions to conventional wisdom are documented. Here, we present one such example by showing that flavi-like viruses can support genome sizes comparable to those observed in the *Nidovirales*. The absence of a detectable exonuclease domain demonstrates that other solutions are available to RNA viruses for overcoming theoretical error thresholds. Future studies may find that the *Nidovirales* and the *Flaviviridae* are not unique in their possession of large genomes. The continued exploration of RNA virus diversity is therefore key to revealing the patterns that shaped the long-term macroevolution of RNA viruses.

## METHODS

### Sample collection

All invertebrate samples analysed here (n = 72) were collected on October 20, 2022, in Chowder Bay, Sydney, Australia. Samples were collected by divers wearing latex gloves and using forceps cleaned with 96% ethanol between sampling events. All samples were placed in RNA/DNA-free cryogenic tubes, which were snap-frozen and stored immediately in liquid Nitrogen. Samples were then transferred to a -80C freezer.

### Sample preparation and metatranscriptome sequencing

Sample tissue was processed individually by flash freezing with liquid nitrogen and homogenising with a mortar and pestle. Total RNA was extracted using the QIAGEN RNeasy Plus Mini kit. The RNA of some samples was pooled for downstream processing. The libraries in which the novel flavi-like virus was found comprised single samples. Sequencing libraries were prepared using the Ribo-Zero Plus library preparation kit and sequenced on the Illumina NovaSeq 6000 platform. In total, 44 libraries were sequenced, including a “blank” negative control comprising a sterile water plus reagent mix.

### Virus discovery

Raw reads were processed by removing sequencing adapters with Trimmomatic v0.38^41^. Reads mapping to rRNA were removed using SortMeRNA v4.3.3^42^ and the SILVA rRNA database (as of Dec. 2023). Contigs were assembled with MEGAHIT v1.2.9^43^ setting the minimum contig length to 200nt. All contigs were screened against the RdRp-scan database^44^ and a custom RNA virus databases using DIAMOND BLASTx v2.0.9^45^ with the setting ultra-sensitive and an e-value cut-off of 1e-5. The custom database included divergent viruses identified in ongoing research projects. Virus candidates (i.e., viruses with a detectable sequence similarity to viruses in one or both of the databases) that were at least 1kb in length were collated for verification. To verify that putative virus sequences were not misassigned host genes, all candidates were screened against the NCBI non-redundant protein database (as of Sept. 2023) using DIAMOND BLAST with an e-value cut-off of 1e-5 and the setting very-sensitive. Virus candidates sharing sequence similarity to host genes were excluded from further analysis. No virus candidates were detected in the negative control.

The sequence of the novel flavi-like virus was translated using Expasy (https://web.expasy.org/translate/) to verify the presence of a complete and uninterrupted open reading frame. Untranslated regions (UTRs) were manually annotated in Geneious Prime v2023.2.1. To further verify that this sequence did not contain known host gene sequences, the polyprotein was screened for sequence and structure homologies using BLASTp and Phyre2^46^.

The abundance of remaining reads (i.e. non-rRNA reads) was estimated using RSEM v.1.3.0^47^ implemented in Trinity v.2.5.1^48^. Raw reads were aligned with bowtie2 v2.3.31^49^. The abundance of the contig encoding the novel flavi-like virus described in this study was calculated by dividing the expected count for that contig by the sum of the expected count for all rRNA reads.

### Genome sequencing coverage

Sequencing coverage was assessed using BBMap v37.98^26^. Calculation of the Q30 score and mapping visualisation was performed by Geneious Prime.

### Host associations

To characterise the host composition of the library, non-rRNA reads and contigs were first screened using KMA v1.3.9a^50^ and CCMetagen v1.1.3^51^. Reads were mapped to the prebuilt KMA database. Next, rRNA reads identified with SortMeRNA v4.3.3^42^ were assembled using MEGAHIT v1.2.9^43^ and screened against the NCBI Blast nt database (as of December 2023).

### Annotation of functional domains

Preliminary annotation of the novel flavi-like virus was performed using InterProScan v2.1 against the CDD, SuperFamily, and NCBIfam databases as implemented in Geneious Prime v20203.2.1. For manual annotation of protein boundaries, transmembrane (TM) domains were predicted using DeepTMHMM v1.0.24^52^. These TM domains were then analysed to determine if they were signal peptidases by querying a 40 amino acid (aa) region encompassing each TM domain using a 15aa sliding window with SignalP v6.0^53^. The potential N-linked glycosylation residues on the polyprotein were identified using NetNGlyc v1.0^54^, with likely N-linked glycan residues considered above the threshold of 0.5. For functional protein domain prediction, the polyprotein sequence was queried against the Pfam v34.0^55^ and CDD v3.20^56^ databases utilising the NCBI CD-search program^57^ with an e-value threshold of 0.1. To examine the presence of a protease, we aligned with NS3Pro of Classical swine fever virus (CSFV), Pangolin pestivirus, and the novel flavi-like virus with MAFFT v7.511^58^ L-INS-I method. CSFV was chosen because it is a pestivirus and the catalytic triad in its protease is well documented. Pangolin pestivirus NS3 shared the highest sequence similarity to that of the putative NS3 region in the novel flavi-like virus according to BLASTp. The putative envelope protein was identified using HHpred^59^, which predicted its presence with 60.1% probability when the first 6000 amino acids of the polyprotein were tested against the PDB_mmCIF70_24_Oct database.

### Protein structure prediction and homology search

We exploited state of the art machine-learning approaches that, for the first time, allow high-confidence *ab initio* protein structure prediction from sequence data alone. The polyprotein sequence of the novel flavi-like virus (12,694 amino acids) was split in to 300 residue blocks each overlapping by 100 residues (resulting in a total of 125 sequence blocks; **Supp. Fig. 6**). The structure of each sequence block was predicted using both AlphaFold^23^ and ESMFold^24^; the average prediction confidence along the length of the polyprotein for either method is provided in **Supp. Fig. 6b**. Whilst the majority of the polyprotein did not yield confident predictions, peaks in confidence corresponding to well-folded domains were produced by both methods. ESMFold performed somewhat better than AlphaFold, likely due to the latter’s requirement for multiple sequence alignments, which cannot be readily assembled for highly divergent sequences, such as those found in this virus. Structure-guided homology searches were performed using the FoldSeek server^25^ and we focussed on hits against experimentally solved structures from the protein database (PDB).

### Phylogenetic analysis

For the family-level virus analysis, all *Flaviviridae* (taxid 11050 and taxid 38144) available on the NCBI Virus were downloaded on Dec. 15, 2022. These data were supplemented with sequences from Mifsud, et al. (2022)^60^, Mifsud, et al. (2024, *in preparation*) and the NCBI nucleotide database (searched using the phrase “flavi[All Fields] OR pesti[All Fields] OR hepaci[All Fields] OR pegi[All Fields] AND viruses[filter]” on Dec. 15, 2022). We then screened the NCBI Virus database for pesti- and pesti-like viruses and large genome flavi-like viruses published in 2023 that were longer than 20kb, and consequently added Macrosiphum euphorbiae virus 1 isolate K01 (YP_009175071, length = 22.8kb) to the data set. Redundant nucleotide sequences were removed using CD-HIT at a cut-off of 95%. Remaining sequences were translated using the Geneious Prime Find ORFs tool (v.2022.0) (www.geneious.com) and trimmed to the NS2/3 and NS5 gene regions, individually. Putative NS5 sequences that lacked the conserved GDD palm motif were excluded. Incomplete genomes were removed manually. The corresponding segments of the two closest protein BLASTp hits to the novel flavi-like virus (Bovine viral diarrhea virus 3 [accession ANW09737] and *Pestivirus brazilense* [accession WEC89329]) were added to each data set. Members of the *Potyviridae* were used as the outgroup for both trees. The final NS2/3 and NS5 data sets contained 425 and 463 sequences, respectively. All data sets were aligned with MAFFT v7.490^58^ using default parameters and the BLOSUM62 scoring matrix (NS2/3 length = 1,048aa; NS5 length = 746aa). Ambiguously aligned regions were removed with a conservation score of 0.15 using trimAl v1.4.1^61^.

For the genus-level analysis, pestivirus and pesti-like virus sequences were realigned using members of the genus *Orthoflavivirus* as the outgroup. Both MAFFT v7.490^58^ and MUSCLE v5.1^62^ alignment algorithms were used. Ambiguously aligned regions were removed at a range of thresholds (**Table 1**) using trimAl v1.4.1^61^.

All phylogenetic trees were inferred using the maximum likelihood method available in IQ-TREE v1.6.12^63^ with 1000 ultrafast bootstraps and substitution model selection restricted to LG models.

To infer the NPDk phylogenetic tree, the 100 BLASTp results to the flavi-associated NDPk domain were downloaded. A sample of NDPk domains from DNA viruses and fungus were added to the data set. Sequences were aligned with MAFFT v7.490^58^, and the ends were manually trimmed to preserve the most conserved region (length = 269 amino acids, gaps inclusive). A maximum likelihood tree was again inferred using IQ-TREE v1.6.12^63^ with 1000 ultrafast bootstraps. The ModelFinder was unrestricted and selected the LG+I+Γ4 substitution model.

## DATA AVAILABILITY

Alignments and trees referred to in this manuscript as well as the nucleotide and amino acid sequences of the flavi-like virus are available on GitHub (https://github.com/mary-petrone/large_flavi).

## ACKNOWLEDGMENTS

This work was funded by a National Health and Medical Research Council (NHMRC) Investigator award and by AIR@InnoHK administered by the Innovation and Technology Commission, Hong Kong Special Administrative Region, China. ECH is supported by a National Health and Medical Research Council Investigator award (GNT2017197). JG is supported by the Wellcome Trust and Royal Society through a Sir Henry Dale Fellowship (107653/Z/15/Z) and the MRC-University of Glasgow Centre for Virus Research core support from the Medical Research Council/UKRI (MC_UU_00034/1). We thank Maxime Cailliau for inspiring the name Maximus pesti-like virus, and we thank Chris Cooney for helping with sample collection.

## AUTHOR CONTRIBUTIONS

M.E.P., E.M.M., and E. C. H. designed the study. E.M.M. collected the samples. M.E.P., J.C.O.M., R.P., and J.G. performed the analyses. M.E.P. and E.C.H. wrote the initial manuscript draft. All authors reviewed and edited the manuscript.

