## Supplementary figures and images for "A 39.8kb flavi-like virus uses a novel strategy for overcoming the RNA virus error threshold"

### Supplementary Figure 1

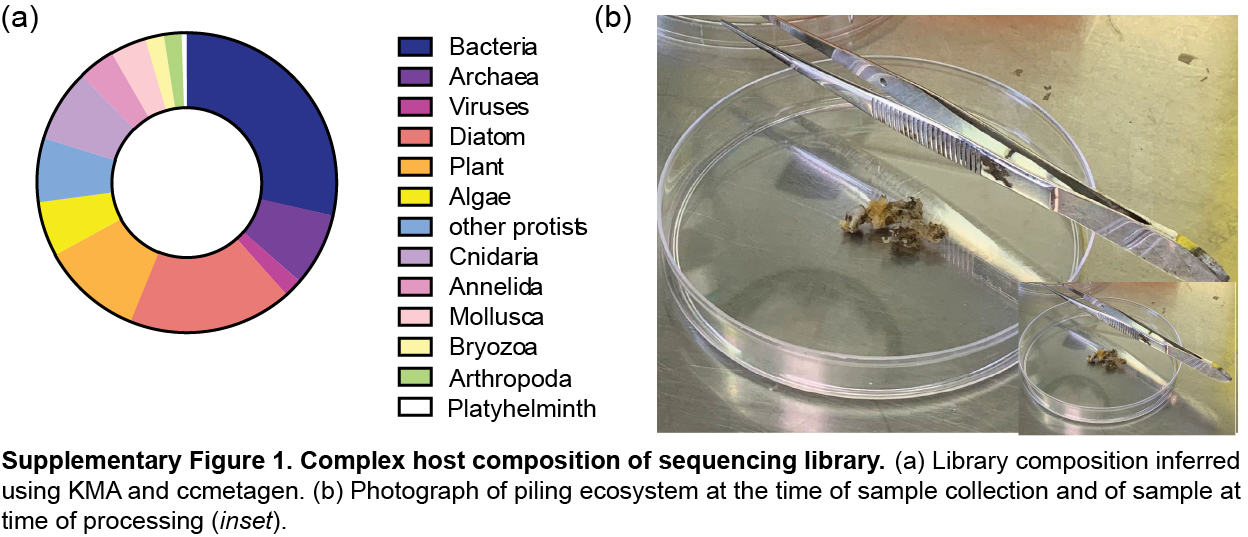
