## Supplementary Figure 2 for "A 39.8kb flavi-like virus uses a novel strategy for overcoming the RNA virus error threshold"

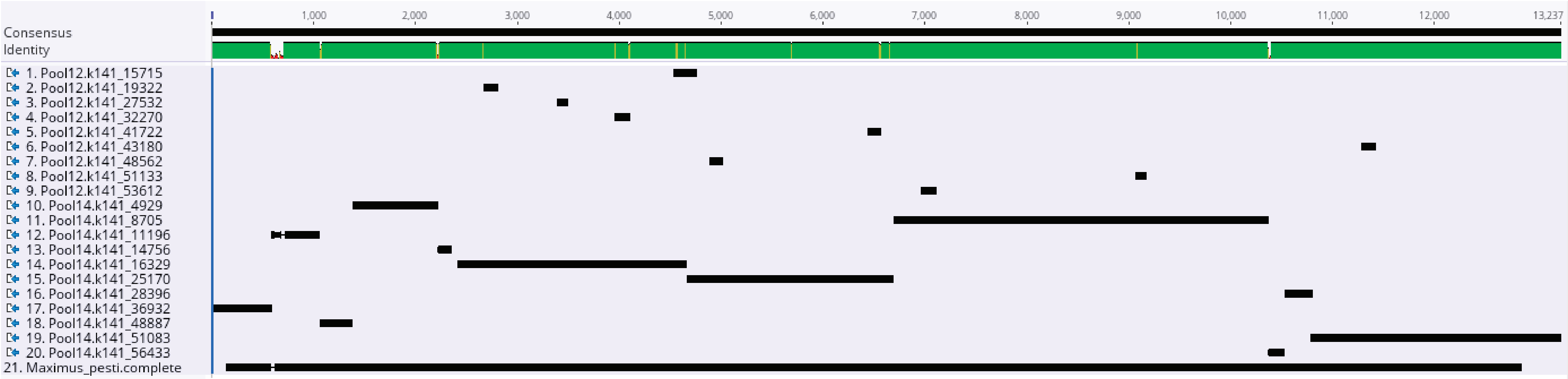


**Supplementary Figure 2. Alignment of Maximus pesti-like virus fragments identified in three sequencing libraries.** Sequences were aligned with CLUSTAL Omega[22] . The alignment was visualised in Geneious Prime. The additional pools are denoted by number (Pool 12 and Pool 14).
