## Supplementary Figure 3 for "A 39.8kb flavi-like virus uses a novel strategy for overcoming the RNA virus error threshold"

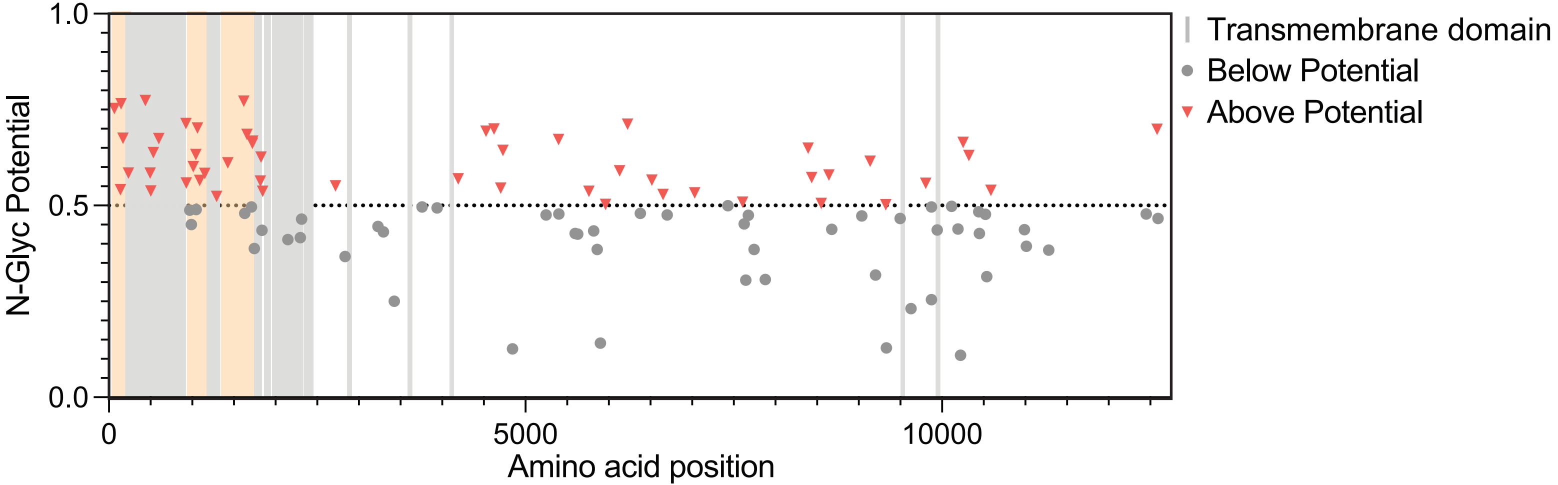


**Supplementary Figure 3. Predicted N-linked glycosylation sites.** Potential N-linked glycosylation residues on the polyprotein were identified using NetNGlyc v1.0[54], with likely N-linked glycan residues considered above the threshold of 0.5. Data were visualised in GraphPad Prism v10.1.0.
