## Supplementary Figure 5 for "A 39.8kb flavi-like virus uses a novel strategy for overcoming the RNA virus error threshold"

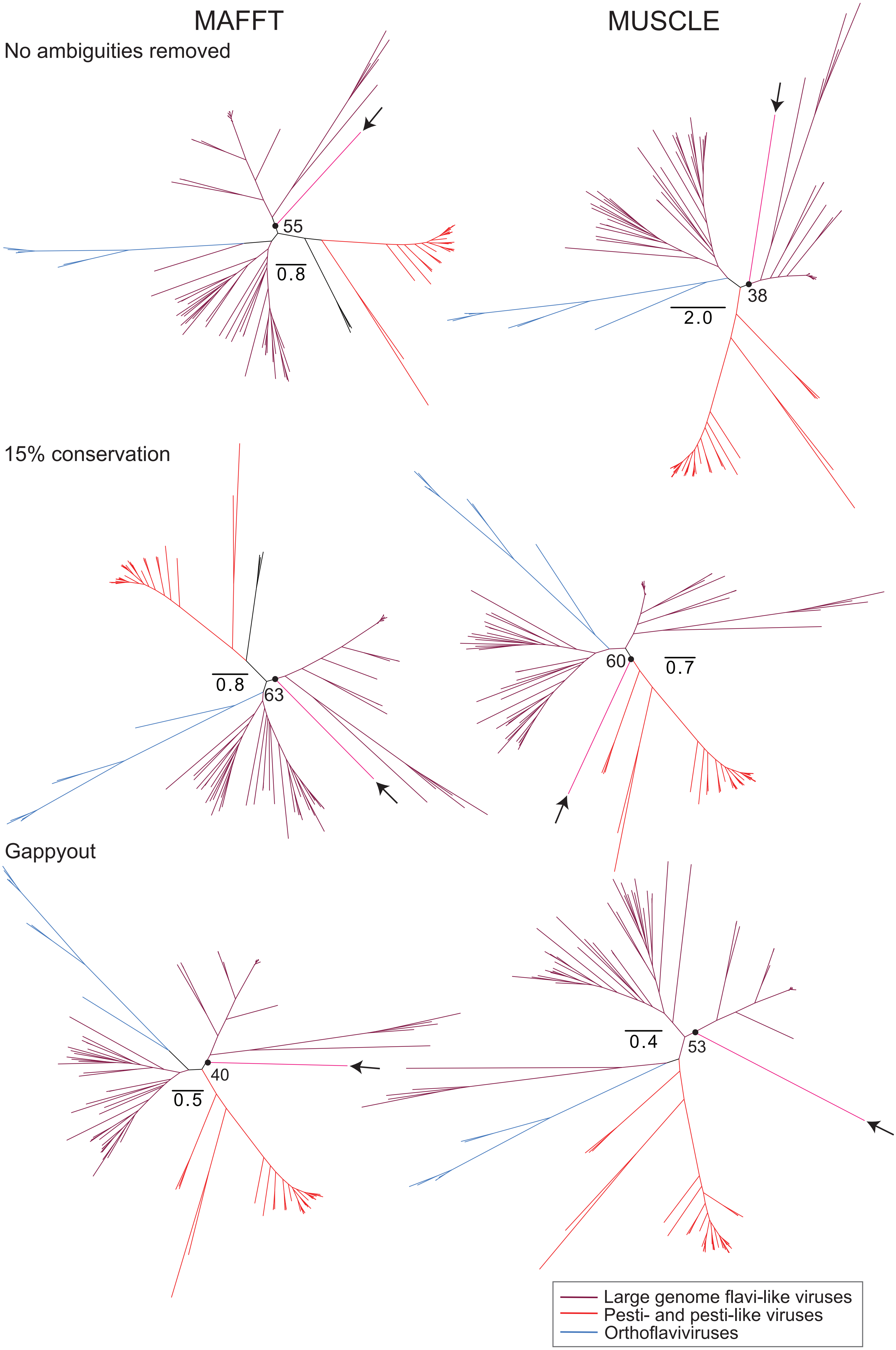


**Supplementary Figure 5.** Unrooted, maximum likelihood phylogenetic trees of selected *Flaviviridae* NS5 inferred with IQ-TREE from six combinations of alignment and trimming methods. Arrows indicate placement of the novel flavi-like virus. Node values indicate ufboot support of this placement.
