## Supplementary Figure 6 for "A 39.8kb flavi-like virus uses a novel strategy for overcoming the RNA virus error threshold"

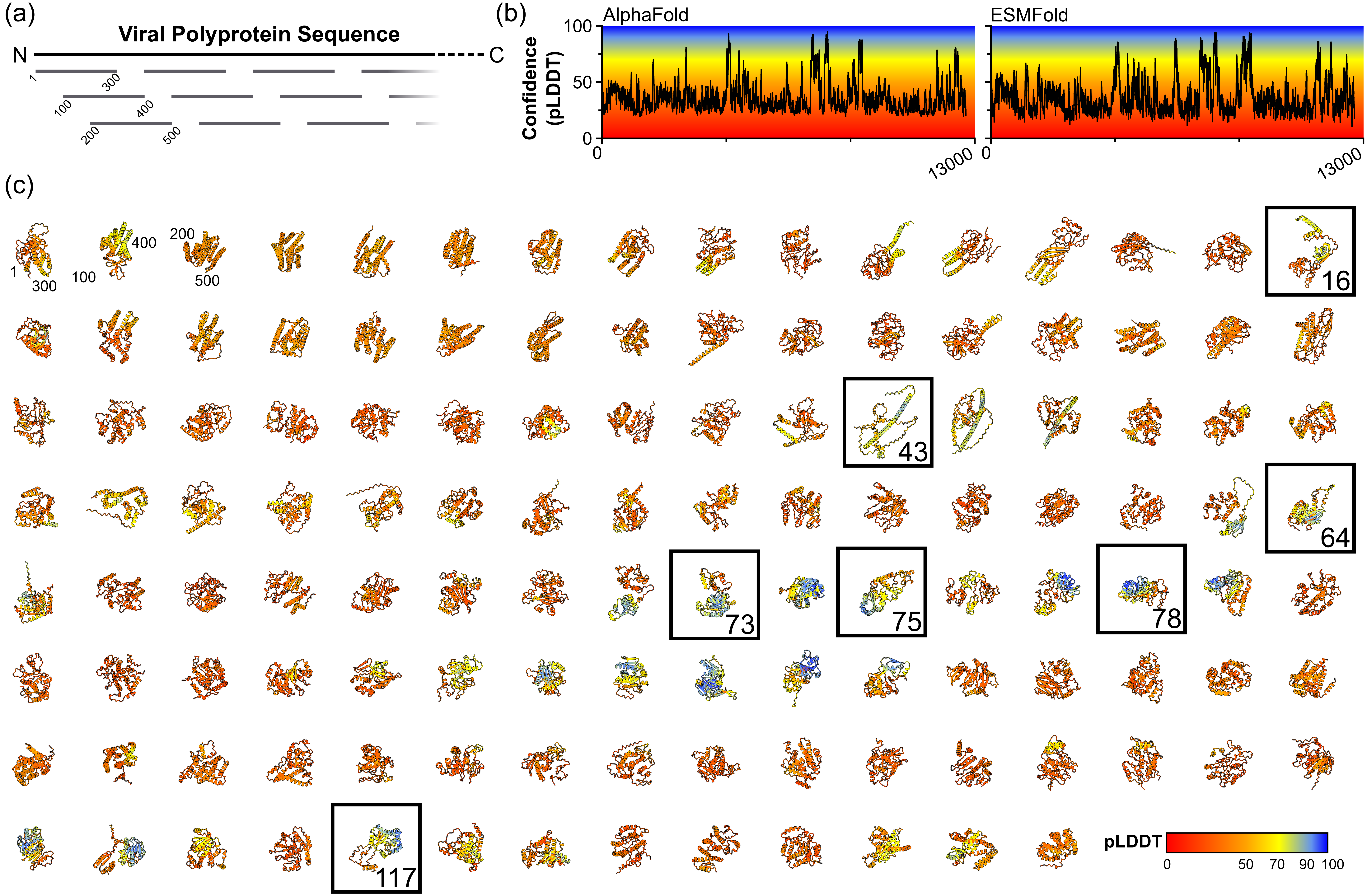


**Supplementary Figure 6. Protein structure prediction and homology search.** (a) Schematic illustrating the overlapping sequence blocks used for structure prediction, residue numberings are shown for the first three blocks. (b) Average prediction confidence plots (predicted Local Distance Difference Test, pLDDT) along the length of the viral polyprotein for structures generated using AlphaFold and ESMFold. (c) ESMFold structure predictions for all 125 sequence blocks color-coded by prediction confidence (pLDDT) as shown in the key and in (b). Blocks that appear in main text figures are outlined and numbered. The N- and C-termini are numbered by residue for the first three blocks, as in (a).
